# Visualizing the Dominant GPCR Coupling of Pathogenic Gαo Mutants in *GNAO1*-Related Disorders

**DOI:** 10.1101/2025.06.19.660437

**Authors:** Yonika A. Larasati, Camille Rabesahala de Meritens, Miriam Stoeber, Vladimir L. Katanaev, Gonzalo P. Solis

## Abstract

Heterozygous mutations in the *GNAO1* gene, which encodes the Gαo subunit of heterotrimeric G proteins, cause a spectrum of rare neurodevelopmental disorders ranging from early-onset epileptic encephalopathy to milder dystonia phenotypes. Disease dominance of Gαo mutants appears to arise from multiple functional disruptions, including impaired guanine nucleotide handling, failure to adopt the active conformation, and neomorphic interactions with Ric8A/B chaperones. In parallel, several Gαo variants have been independently reported to dominantly engage G protein-coupled receptors (GPCRs) or to sequester Gβγ, two mechanistically distinct behaviors. These conclusions were inferred from different indirect biosensor assays, likely contributing to the apparent contradiction in their proposed mechanisms. To overcome this, we developed a split-YFP-based bimolecular fluorescence complementation (BiFC) assay to visualize receptor-Go protein complexes at the plasma membrane. Using this system, we found that severe Gαo variants (S47G, G203R, R209C, E246K) fail to disengage from activated Gi/o-coupled GPCRs, thereby preventing downstream receptor phosphorylation and endocytosis. By contrast, milder dystonia-linked mutants (C215Y and T241_N242insPQ) showed near-normal receptor internalization and only minor phosphorylation defects. These findings establish dominant GPCR coupling as a molecular hallmark of severe *GNAO1*-related disorders and point to split-YFP BiFC as a robust platform for probing mutant G protein behavior in genetic disease.

## Introduction

G protein-coupled receptors (GPCRs) represent the largest and most diverse family of membrane receptors in metazoans^1–3^. From neurotransmission and hormonal signaling to sensory perception and immune responses, GPCRs are essential for proper animal development and physiology^4, 5^. Signal transduction by GPCRs is mediated by heterotrimeric G proteins, composed of three subunits: α, β, and γ, with the Gα subunit conferring receptor specificity^6, 7^. In the inactive state, Gα is bound to GDP and associates tightly with the Gβγ dimer. Upon ligand binding, the GPCR undergoes a conformational change that catalyzes the exchange of GDP for GTP on Gα. This process activates the heterotrimeric G protein, causing Gα-GTP and Gβγ to dissociate and signal independently to downstream effectors.

The signal is terminated when Gα hydrolyzes GTP to GDP through its intrinsic GTPase activity and associates with Gβγ, reforming the heterotrimer^8, 9^. Given their central role in GPCR signaling, it is perhaps unsurprising that mutations affecting heterotrimeric G protein subunits have been implicated in a variety of human diseases.

Historically, gain-of-function mutations in genes encoding Gα subunits have drawn most of the attention due to their role as drivers in various cancers^10, 11^. These mutations produce constitutively active Gα proteins, resulting in persistent stimulation of oncogenic signaling pathways. More recently, however, a growing number of rare genetic disorders have been attributed to alternative mutations in Gα subunits^12–17^. Among these, mutations in *GNAO1* – the gene encoding for Gαo – have been implicated in a broad spectrum of neurodevelopmental disorders, collectively referred to as *GNAO1*-related disorders^13, 18–20^. The most severe conditions were originally classified as Developmental and Epileptic Encephalopathy 17 (DEE17; OMIM #615473) and Neurodevelopmental Disorder with Involuntary Movements (NEDIM; OMIM #617493)^21, 22^. In recent years, *GNAO1* mutations have also been associated with milder conditions, including autism and intellectual disability, parkinsonism, and dystonia^23–25^. More than 250 patients have been reported worldwide, the vast majority carrying missense mutations, with only a few cases of deletions, frameshifts, duplications, short indels, and splice-site mutations described^18^. Treatment responses among *GNAO1* patients are highly variable and typically poor, particularly in cases involving epilepsy and/or movement disorders. This underscores the urgent need for more effective therapies for these severe conditions.

At the molecular level, Gαo mutants associated with severe phenotypes display a set of interconnected pathogenic mechanisms, including severely impaired guanine nucleotide handling, misfolding in the GTP-bound active state, and neomorphic binding to Ric8A/B chaperones, all of which converge to perturb the broader GPCR signaling network^26–30^. By contrast, variants on the milder end of the *GNAO1*-related disorder spectrum exhibited only mild biochemical defects, with some mutants lacking the neomorphic property entirely, while others retained trapping of Ric8A but not Ric8B^24, 25, 28, 31^. Furthermore, the most severe DEE17 Gαo variants tend to lose plasma membrane (PM) localization and Gβγ binding, whereas mutants linked to NEDIM or milder phenotypes retain near-normal association with both PM and Gβγ^26–28^.

Another emerging feature of several pathogenic *GNAO1* mutations is their apparent dominant coupling to GPCRs. A study using a bioluminescence resonance energy transfer (BRET)-based assay to measures free Gβγ during GPCR stimulation showed that clinically severe Gαo variants partially inhibit activation of the wild-type protein, acting in a dominant-negative (DN) manner^26^. These Gαo mutants displayed poor disengagement from Gβγ despite normal (or even enhanced) coupling to activated GPCRs – the latter determined indirectly by a split-NanoLuc assay measuring Gβγ proximity to the receptor. Based on this, the authors proposed that the DN effect arises from nonproductive interactions between the mutant Gαo and GPCRs. Along the same lines, we found that many clinically severe Gαo mutants exhibit normal or even enhanced GPCR coupling despite poor binding to Gβγ, using a different BRET-based assay that directly monitors GPCR engagement by Gα^28^. This dominant coupling mechanism was recently confirmed for the epileptic Gαo K46E variant, which forms a nonproductive complex with both Gβγ and the activated GPCR, as revealed by cryo-electron microscopy^32^. By contrast, an earlier study using yet another BRET-based assay that directly measures Gβγ disengagement from Gα described an alternative DN mechanism for some of the same Gαo mutants: sequestration of Gβγ, due to impaired adoption of the active conformation required for their dissociation, without sequestering the GPCR^27^.

Thus, dominant GPCR coupling by pathogenic Gαo mutants can currently only be inferred using a combination of optical biosensors, leading to apparent mechanistic contradictions. To resolve this, we developed a novel assay to directly visualize the dominant GPCR coupling behavior of Gαo variants. We employed bimolecular fluorescence complementation (BiFC), a method based on the structural reassembly of two non-fluorescent N- and C-terminal fragments of a fluorescent protein into a functional fluorophore when both fragments are in close proximity^33^. Using the split-YFP-based BiFC approach, we show that *GNAO1* mutations associated with the most severe phenotypes result in dominant GPCR coupling by the mutant Gαo, likely caused by a defective receptor-Gα uncoupling.

## Results

### Dominant GPCR coupling by pathogenic Gαo mutants blocks receptor endocytosis

To determine whether pathogenic Gαo mutants indeed engage in dominant coupling with GPCRs, we sought to develop a single assay capable of visualizing this process. However, microscopic visualization of plasma membrane (PM)-localized GPCRs can be challenging, as exogenous expression often leads to their accumulation in the endoplasmic reticulum and Golgi apparatus, due to saturation of the cellular folding and trafficking machinery^34, 35^.

Consistent with this, expression of an N-terminal GFP fusion of the M_2_ muscarinic acetylcholine receptor (GFP-M_2_R) in HEK293T cells results in prominent intracellular retention (Fig. S1A). We selected M_2_R for this study because it is a prototypical Gi/o-coupled GPCR^36, 37^ and because we previously observed that severe Gαo mutants exhibit enhanced coupling to this receptor^28^, consistent with dominant GPCR engagement.

To overcome this limitation, we employed a split-YFP-based BiFC assay (hereafter split-YFP), previously used to visualize the PM interaction between the chemokine GPCR CCR7 and the Src tyrosine kinase^38^. Like Src, Gα subunits associate with the membrane via lipid modifications^39, 40^, which target Gαo to the PM and Golgi in HEK293T cells (Fig. S1B)^41^.

Based on these observations, we reasoned that a split-YFP assay combining M_2_R and Gαo would enable selective visualization of the PM-localized receptor pool. We further speculated that the irreversible nature of the split-YFP complementation would bias M_2_R toward coupling with Gαo over endogenous Gα subunits, thereby enhancing both the specificity and sensitivity of the assay.

We first generated M_2_R and Gαo constructs carrying each of the YFP fragments – N-terminal (YN) or C-terminal (YC) – fusing them to the intracellular C-terminus of the receptor and internally at Gly92 in Gαo (Fig 1A). One of the two possible combinations – M_2_R-YN and Gαo-YC (M_2_R-YFP-Gαo; Fig. 1B) – produced a much stronger fluorescence signal than the opposite configuration (not shown), confirming that the orientation of the YFP halves is critical for efficient complementation^42^. As speculated, the M_2_R-YFP-Gαo complex was strongly visualized at the PM of HEK293T cells, with only a few intracellular structures visible (Fig. 1B). Notably, acetylcholine (ACh) stimulation for 10 min led to the prominent appearance of vesicles/endosomes and a marked reduction in PM signal, suggesting endocytosis of the M_2_R-YFP-Gαo complex (Fig. 1C). Although Gαo typically remains at PM following GPCR stimulation^41, 43^, the irreversibility of split-YFP complementation likely drives its co-internalization with the receptor^33^.

**Figure 1.**
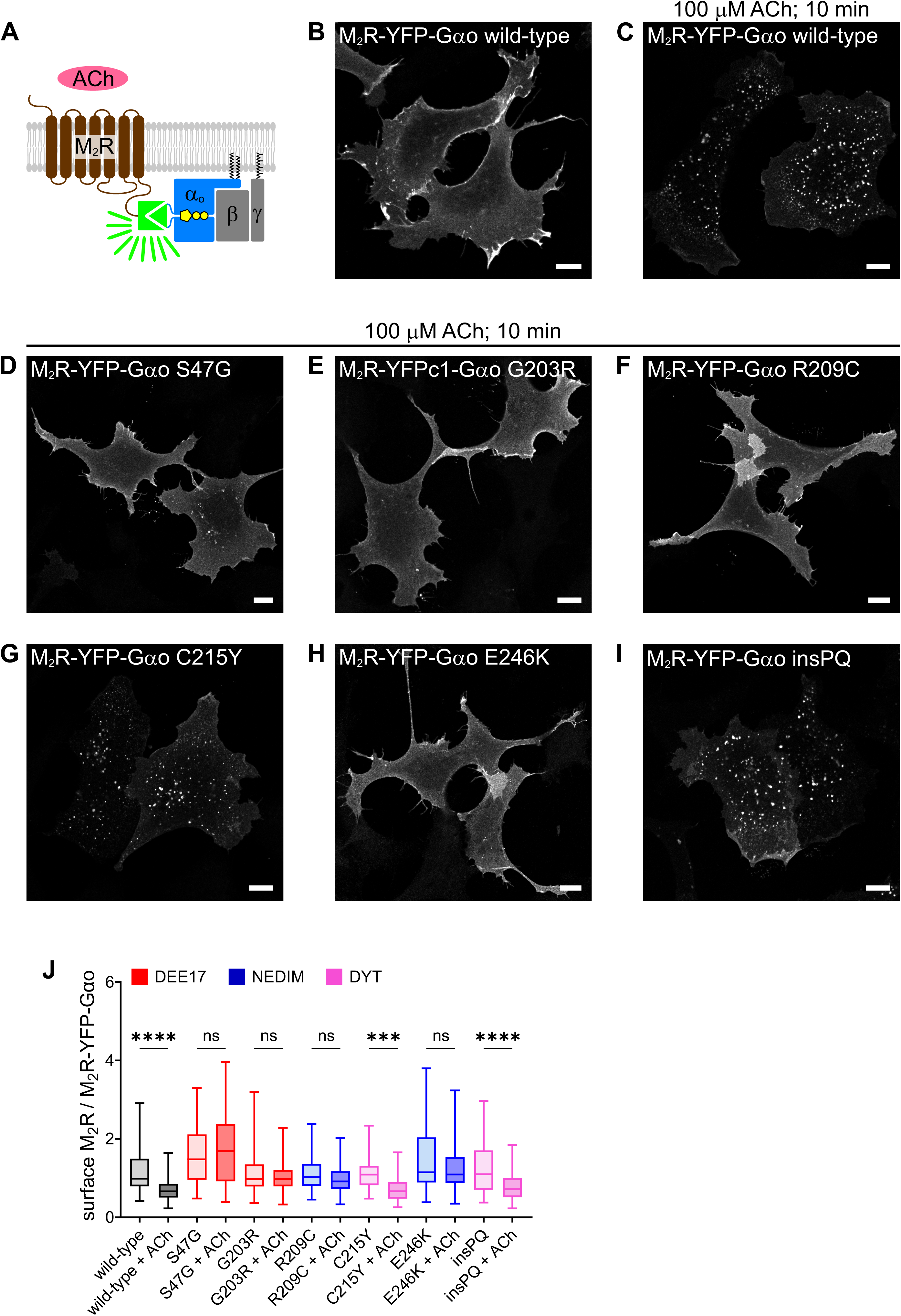
Clinically severe Gαo mutants disrupt M_2_R endocytosis. (**A**) Illustration of the split-YFP assay applied to M_2_R and heterotrimeric Gαoβγ. (**B**,**C**) Representative confocal images of HEK293T cells expressing the M_2_R-YFP-Gαo complex at steady state (**B**) and after 10 min of acetylcholine (ACh) stimulation (**C**). (**D**-**I**) Confocal images of ACh-stimulated HEK293T cells expressing the M_2_R-YFP-Gαo complex containing the Gαo mutants S47G (**D**), G203R (**E**), R209C (**F**), C215Y (**G**), E246K (**H**), and insPQ (**I**). Scale bars, 10 µm. (**J**) Quantification of ACh-mediated M_2_R internalization, expressed as the surface level of M_2_R normalized to total M_2_R-YFP-Gαo signal. Gαo mutant associations with DEE17, NEDIM and DYT phenotypes are color-coded. Box plot indicate median (middle line), 25th, 75th percentile (box), and lowest, highest value (whiskers); two-three independent experiments (wild-type, *n* = 78; wild-type +ACh, *n* = 85; S47G, *n* = 55; S47G + ACh, *n* = 51; G203R, *n* = 59; G203R + ACh, *n* = 69; R209C, *n* = 78; R209C + ACh, *n* = 76; C215Y, *n* = 59; C215Y + ACh, *n* = 74; E246K, *n* = 63; E264K + ACh, *n* = 65; insPQ, *n* = 54; insPQ + ACh, *n* = 63). Statistical analysis was performed using one-way ANOVA followed by Šídák’s multiple comparisons test; \*\*\**p* < 0.001, \*\*\*\**p* < 0.0001, and ns: not significant.

Next, we performed the split-YFP assay for M_2_R and several pathogenic Gαo mutants. We selected three of the most recurrent Gαo variants – G203R, R209C, and E246K – that may act as DN, either through nonproductive GPCR coupling^26, 28^ or via Gβγ sequestration^27^. We also included the S47G mutant, which shows normal to high GPCR coupling but poor Gβγ dissociation^26, 28^, and has alternatively been described as non-functional^27^. Among these, Gαo S47G and G203R are associated with the most severe DEE17 phenotype, while R209C and E246K are linked to NEDIM. Two Gαo mutants associated with a milder dystonia (DYT) phenotype were also analyzed: C215Y and the recurrent splice-site variant c.724-8G>A, which results in an in-frame insertion of two additional residues (Pro-Gln) at position T241 (T241_N242insPQ; hereafter insPQ)^23, 28, 31, 44^.

As shown in Figure 1, split-YFP complementation was achieved for all pathogenic Gαo mutants. However, ACh stimulation revealed striking differences, with DEE17 and NEDIM variants strongly impairing internalization of the M_2_R-YFP-Gαo complex and DYT mutants showing no apparent defects (Fig. 1D-I). To quantify endocytosis, we analyzed the PM localization of M_2_R-YN before and after ACh stimulation using immunostaining – under non-permeabilizing conditions – against an extracellular HA-epitope at the N-terminus of the construct (Fig. S2A-G). The extracellular HA-signal was normalized to the intrinsic fluorescence of the M_2_R-YFP-Gαo complex to account for variability in expression levels across cell populations. As expected, a significant reduction of the relative PM pool of M_2_R was observed only for wild-type Gαo and the mild DYT variants C215Y and insPQ (Fig. 1J). For these Gαo mutants, we noticed that the HA-derived M_2_R signal was strongly reduced upon ACh stimulation in some cells, but not in others – particularly those with higher expression levels (Fig. S2A-G). This variability may reflect saturation of the receptor endocytic pathway due to overexpression^45^.

We also noticed that the overall YFP signal in cells expressing M_2_R-YFP-Gαo complexes was reduced for some of the pathogenic variants compared to wild-type. Quantification revealed that the DEE17 S47G, NEDIM E246K, and DYT insPQ variants formed significantly lower levels of M_2_R-YFP-Gαo complexes (Fig. S2H). This is likely the outcome of reduced expression of the corresponding Gαo-YC constructs (Fig. S2I and J), consistent with the known tendency of many pathogenic Gαo mutants to express at lower levels^27, 28^. These expression differences, however, do not account for the defective internalization observed for DEE17 and NEDIM variants, as the DYT insPQ variant – despite a similar reduction in M_2_R-YFP-Gαo formation – underwent robust internalization.

The most straightforward explanation for the failure of M_2_R-YFP-Gαo complexes formed by DEE17/NEDIM variants to undergo endocytosis is a defect in receptor-Gαo uncoupling, which prevents progression toward receptor internalization. However, we cannot exclude the possibility that these severe Gαo mutants may impair endocytosis more broadly. To address this, we examined the internalization of an unrelated receptor, epidermal growth factor receptor (EGFR), using fluorescently labelled EGF-Rhodamine as readout^46^. EGF-Rhodamine uptake in HEK293T cells expressing the M_2_R-YFP-Gαo complex was comparable to that in non-transfected neighboring cells (Fig. S3A-G). Furthermore, quantification of internalized EGF-Rhodamine in cells expressing M_2_R-YFP-Gαo revealed no significant differences among the various Gαo variants (Fig. S3H). These results suggest that expression of DEE17/NEDIM Gαo mutants does not broadly interfere with receptor endocytosis.

Next, we tested the specificity of the split-YFP assay using different GPCRs. We selected the µ-opioid receptor (MOR; Fig. S4A), which is known to engage Gαo, and the β_2_- adrenoceptor (β_2_AR; Fig. S5A), which signals primarily through Gαs^47, 48^. Notably, split-YFP complementation was effectively achieved between wild-type Gαo-YC and either MOR-YN or β_2_AR-YN, with MOR-YFP-Gαo (Fig. S4B) and β_2_AR-YFP-Gαo (Fig. S5B) complexes localized primarily at the PM of HEK293T cells. These results indicate that split-YFP complementation with Gαo is not selective for Gi/o-coupled receptors. Recapitulating the pattern observed with M_2_R, stimulation with fentanyl for 10 min induced internalization of the wild-type MOR-YFP-Gαo complex into vesicles/endosomes (Fig. S4C), a pattern also observed for the DYT variants, but not for the mutants associated with DEE17/NEDIM (Fig. S4D-I). Conversely, isoproterenol treatment induced the endocytosis of the β_2_AR-YFP-Gαo complex formed by the wild-type protein and all pathogenic variants alike (Fig. S5C-I), suggesting that the dominant GPCR coupling of Gαo mutants is specific for Gi/o-coupled receptors. This finding further implies that split-YFP complementation between β_2_AR and Gαo does not block the ability of the receptor to couple with endogenous G proteins.

### ACh-mediated M_2_R phosphorylation is blocked by dominant Gαo variants in the split-YFP assay

Following the dissociation of G proteins from activated GPCRs, a cascade of molecular events is initiated. GPCR kinases (GRKs) are recruited to the receptors, where they bind to the same intracellular pocket previously occupied by the Gα subunit^49^. GRK recruitment enables phosphorylation of multiple residues within the intracellular loops and/or C-terminus of the GPCR. This promotes the subsequent recruitment of β-arrestins, which act as scaffolds for the clathrin-mediated endocytic machinery, ultimately driving receptor internalization^50^.

Our findings support the notion that the most severe pathogenic Gαo variants fail to disengage from activated GPCRs, remaining dominantly coupled and thereby preventing GRK recruitment and subsequent receptor phosphorylation. To test this prediction, we analyzed M_2_R phosphorylation using the split-YFP assay. We used a non-pathogenic Gαo construct carrying a four-alanine insertion in the C-terminal α5-helix (ins4A) as control, which is known to abolish G protein release from activated receptors^51, 52^. As expected, Gαo ins4A prevented ligand-mediated internalization of both M_2_R (Fig. S6A-C) and MOR (Fig. S6D-F) in the split-YFP assay.

GRK-mediated phosphorylation of M_2_R targets several Ser/Thr residues in the third intracellular loop (Fig. 2A)^53^. We immunostained HEK293T cells expressing the M_2_R-YFP- Gαo complex using a specific antibody that recognizes dual phosphorylation at Thr307 and Ser309 (pT307/pS309). In unstimulated cells, the anti-pT307/pS309 signal was minimal in both transfected and non-transfected cells (Fig. 2B). However, following 7 min of ACh stimulation, robust anti-pT307/pS309 staining appeared and closely colocalized with endocytic M_2_R-YFP-Gαo–positive vesicles (Fig. 2C). Neighboring untransfected cells showed no increase in the pT307/pS309 signal after stimulation, consistent with the absence of endogenous M_2_R expression in the parental HEK293 line^54^.

**Figure 2.**
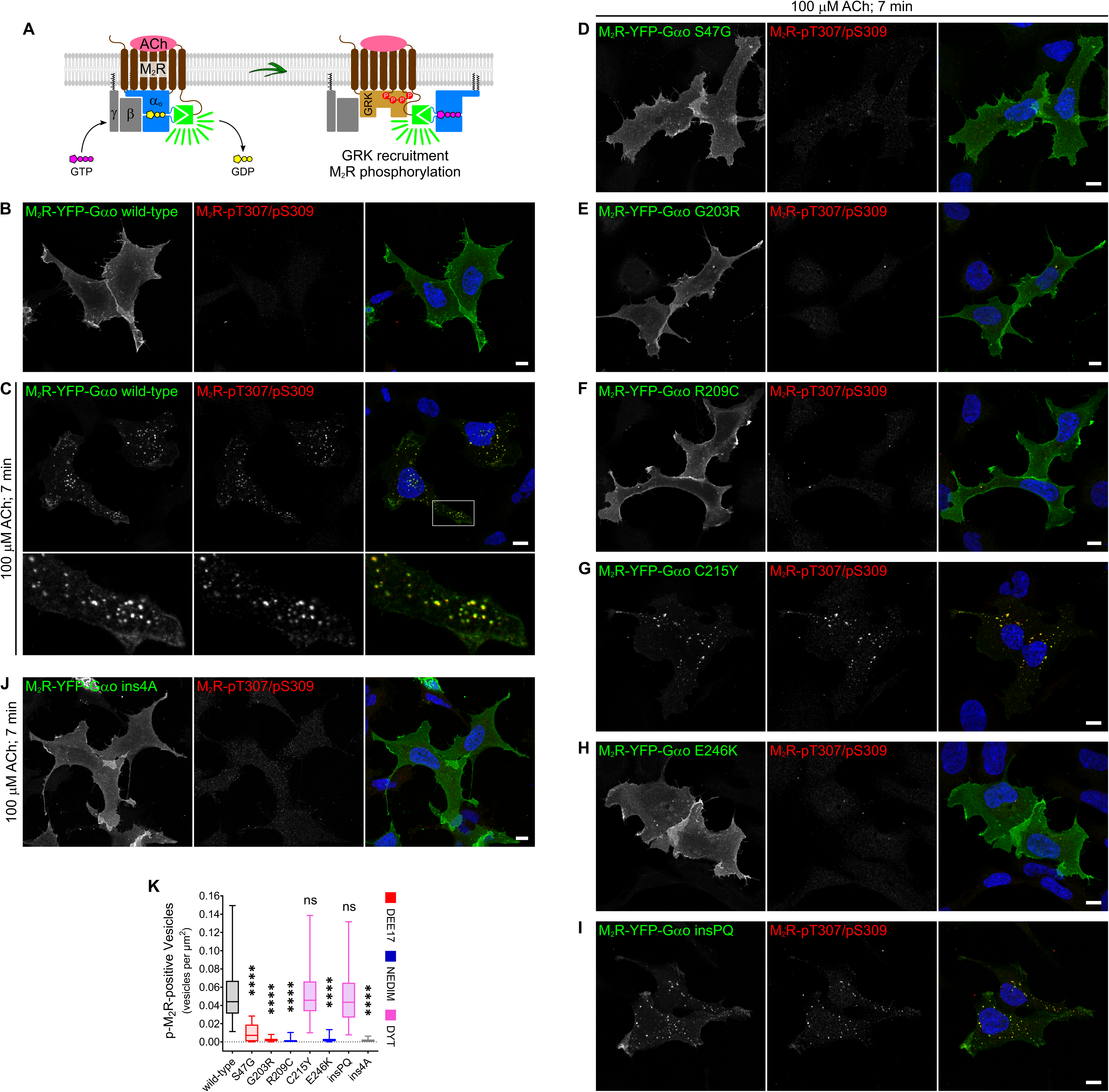
Inhibition of ACh-mediated phosphorylation of M_2_R by pathogenic Gαo variants. (**A**) Schematic representation of the molecular events following ACh stimulation that lead to M_2_R phosphorylation, as illustrated using the split-YFP assay. (**B**,**C**) Representative confocal images of HEK293T cells expressing the M_2_R-YFP-Gαo complex at steady state (**B**) and after 7 min of acetylcholine (ACh) stimulation (**C**). Cells were immunostained with a specific antibody recognizing dual phosphorylation at T307 and S309 of M_2_R (M_2_R-pT307/pS309), and counterstained with DAPI to visualize nuclei. A boxed region in (**C**) is shown at higher magnification in the lower panels. (**D**-**J**) ACh-stimulated HEK293T cells expressing the M_2_R- YFP-Gαo complex with the Gαo mutants S47G (**D**), G203R (**E**), R209C (**F**), C215Y (**G**), E246K (**H**), insPQ (**I**), and the control ins4A (**J**). Cells were prepared as in (**C**). Scale bars, 10 µm. (**K**) Quantification of ACh-mediated endocytic events, expressed as the number of vesicle-like structures positive for both M_2_R-YFP-Gαo and anti-pT307/pS309 staining, normalized to cell area. Associations of Gαo mutants with DEE17, NEDIM, and DYT phenotypes are color-coded. Box plot indicate median (middle line), 25th, 75th percentile (box), and lowest, highest value (whiskers); two-three independent experiments (wild-type, *n* = 46; S47G, *n* = 43; G203R, *n* = 46; R209C, *n* = 44; C215Y, *n* = 48; E246K, *n* = 47; insPQ, *n* = 50; ins4A, *n* = 46). Statistical analysis was performed using one-way ANOVA followed by Dunnett’s multiple comparisons test; \*\*\*\**p* < 0.0001 and ns: not significant.

Notably, the Gαo variants associated with the severe DEE17/NEDIM phenotypes exhibited weak pT307/pS309 staining following ACh stimulation (Fig. 2D-F and H). Since the ins4A construct produced an equivalent effect (Fig. 2J), this result supports the notion that Gαo mutants dominantly couple to M_2_R, thereby preventing GRK recruitment. In contrast, the DYT-associated mutants retained robust phospho-M_2_R staining, similar to wild-type Gαo (Fig. 2G and I). To quantify endocytic events, we counted the number of vesicle-like structures positive for both M_2_R-YFP-Gαo and pT307/pS309, and normalized to the cell area (Fig. 2K). A marked reduction in phospho-M_2_R-YFP-Gαo–positive vesicles was observed for the DEE17/NEDIM variants and ins4A, whereas DYT mutants exhibited values comparable to wild-type.

To confirm the defects in ACh-mediated M_2_R phosphorylation, we took advantage of an anti-GFP nanobody^55^ that enables specific immunoprecipitation (IP) of the reconstituted split-YFP, but not the individual YN and YC fragments (Fig. S7). In Western blots following IPs, four distinct protein species can be detected using M2R- and Gαo-specific antibodies: the M_2_R-YN and Gαo-YC monomers, a monomeric M_2_R-YFP-Gαo complex, and a higher molecular weight species, possibly corresponding to a M_2_R-YFP-Gαo dimer (Fig. S7). This likely reflects the known tendency of M_2_R to form dimers and higher-order oligomers^56^.

Corroborating the cell imaging data, IP of split-YFP complexes containing wild-type Gαo showed no detectable anti-pT307/pS309 signal for M_2_R at steady state, whereas strong signals emerged following 5 min of ACh stimulation in three species: M_2_R-YN, and the monomeric and dimeric forms of the M_2_R-YFP-Gαo complex (Fig. 3A). In contrast, M_2_R- YFP-Gαo complexes formed by the most severe DEE17/NEDIM mutants exhibited a marked reduction in anti-pT307/pS309 reactivity. Unexpectedly, both DYT-associated Gαo variants showed a modest reduction in M_2_R phosphorylation. Quantification revealed a 60-85% reduction for the DEE17/NEDIM variants, and over 90% for Gαo ins4A (Fig. 3B). DYT mutants, on the other hand, showed a significant but smaller 20-25% decrease, suggesting that these variants partially impair GRK recruitment – although not to a degree that prevents receptor endocytosis.

**Figure 3.**
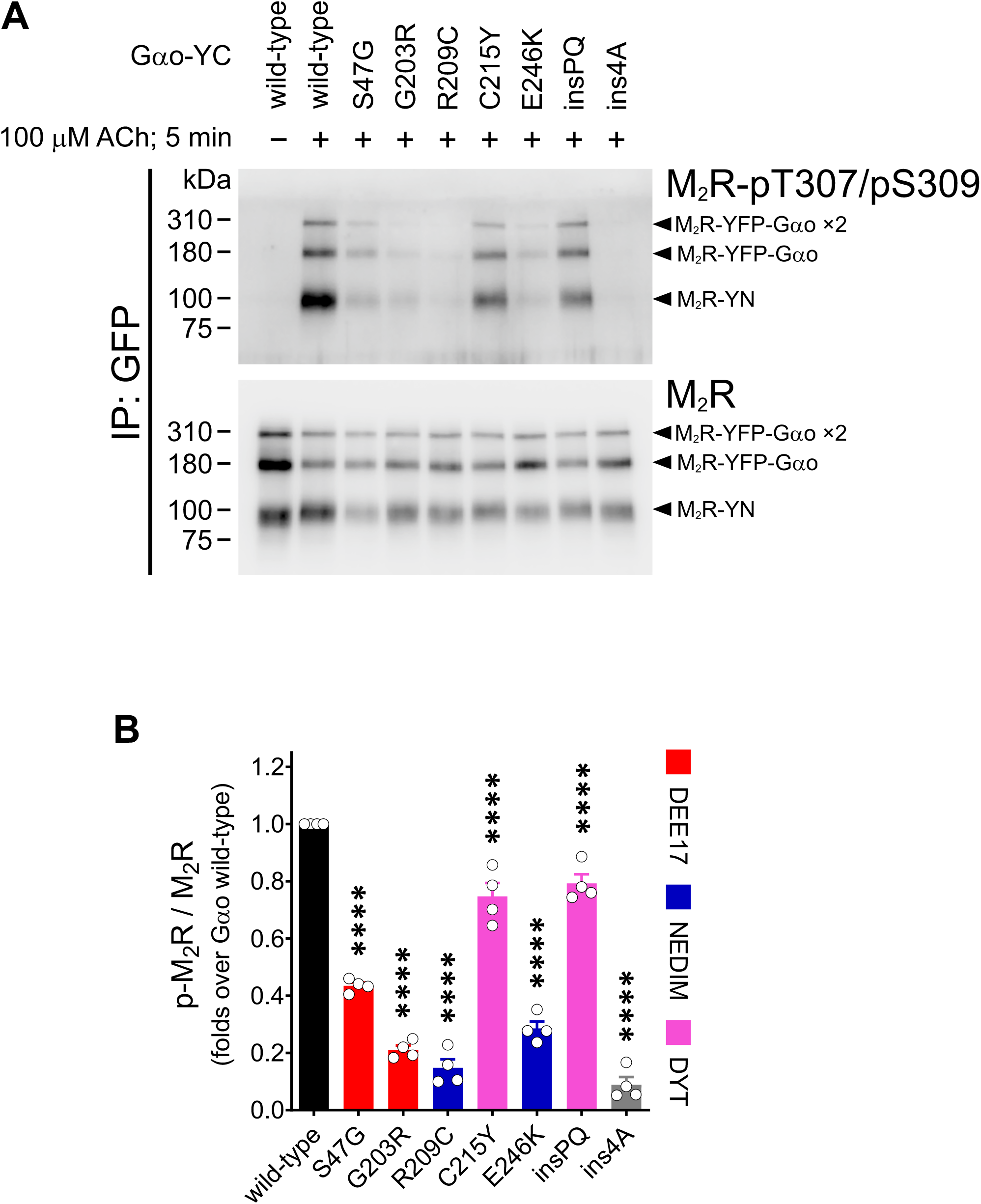
Immunoprecipitation of the M_2_R-YFP-Gαo complex confirms impaired phosphorylation by pathogenic Gαo variants. (**A**) HEK293T cells expressing the split-YFP constructs M_2_R-YN and Gαo-YC (wild-type, pathogenic mutants, and the ins4A control) were stimulated with 100 µM acetylcholine (ACh) for 5 min and subjected to immunoprecipitation (IP) using an anti-GFP nanobody. Western blotting and immunodetection were performed with antibodies against phosphorylated (pT307/pS309) and total M_2_R. Arrowheads indicate M_2_R-YN, and the monomeric and dimeric forms of the M_2_R-YFP-Gαo complex. (**B**) Quantification of M_2_R phosphorylation levels relative to total receptor (*n* = 4). Gαo mutant associations with DEE17, NEDIM, and DYT phenotypes are color-coded. Data represent mean ± SEM. Statistical analysis was performed using one-way ANOVA followed by Dunnett’s multiple comparisons test; \*\*\*\**p* < 0.0001.

### Untagged Gαo mutants recapitulate M_2_R endocytosis and phosphorylation defects

While the split-YFP assay provides a powerful tool to visualize these interactions, it relies on internally tagged Gαo constructs, which could theoretically introduce artifacts not present in native, untagged proteins. To address this, we next tested whether the observed effects also occur with untagged Gαo variants.

We first employed the split-YFP assay for M_2_R and Gγ3 (Fig. 4A), and found that the combination of M_2_R-YC and YN-Gγ3 produced a much stronger fluorescence signal than the reverse configuration (not shown). To test dominant GPCR coupling and receptor endocytosis, we expressed different construct sets in two separate HEK293T cell populations – M_2_R-YC and YN-Gγ3 with or without untagged Gαo – and then co-cultured them. After 7 min of ACh treatment, cells were immunostained for Gαo and pT307/pS309.

**Figure 4.**
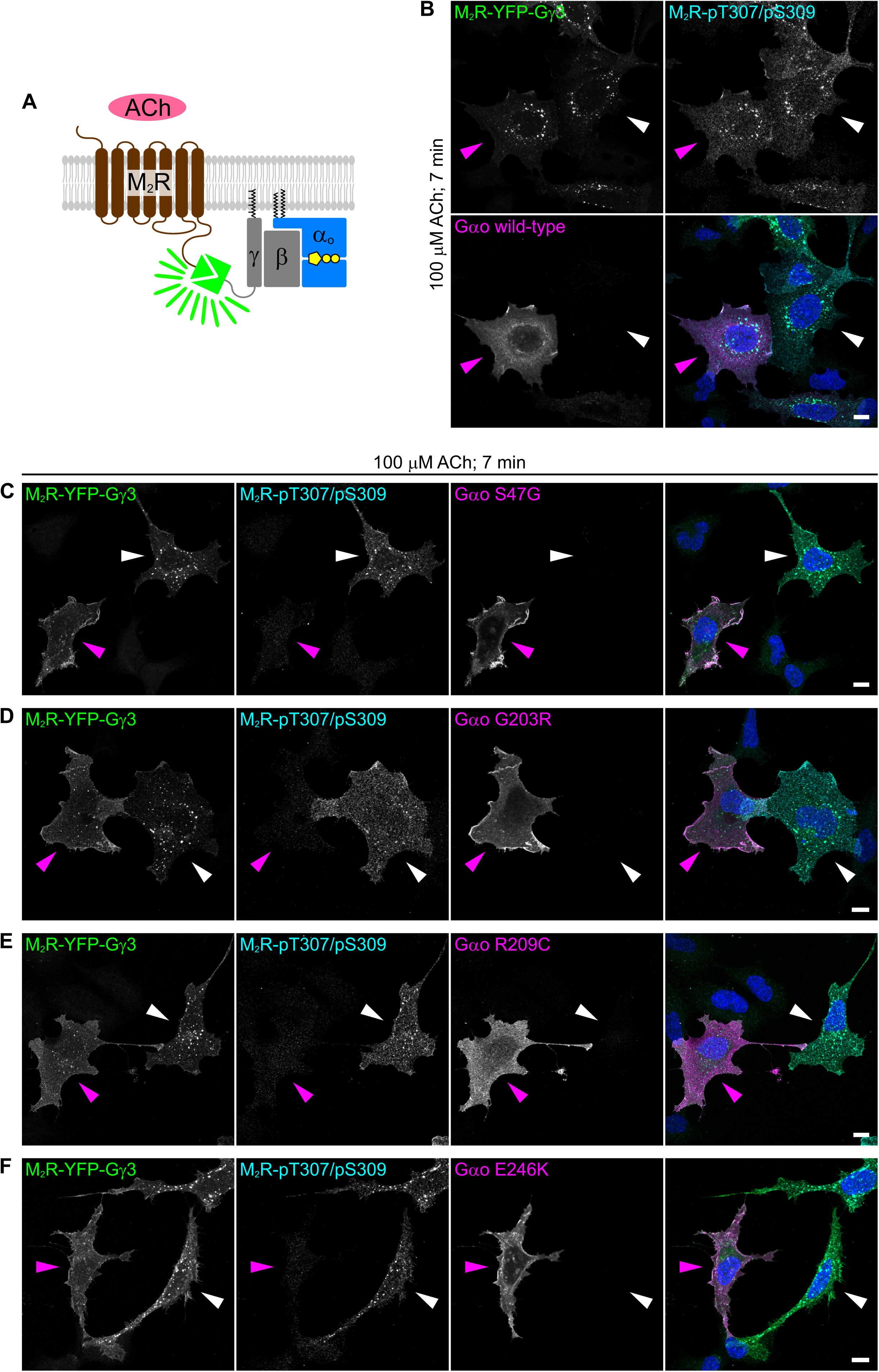
Split-YFP assay with M_2_R and Gγ3 recapitulates phosphorylation and endocytosis defects caused by Gαo mutants. (**A**) Depiction of the split-YFP assay applied to M_2_R and Gγ3. (**B**-**F**) Confocal images of co- culture of HEK293T cells expressing the M_2_R-YFP-Gγ3 complex, with or without co-expression of Gαo wild-type (**B**), S47G (**C**), G203R (**D**), R209C (**E**), and E246K (**F**). Cells were stimulated with 100 µM acetylcholine (ACh) for 7 min, then immunostained with anti- pT307/pS309, anti-Gαo, and counterstained with DAPI to visualize nuclei. Magenta and white arrowheads indicate Gαo-expressing and non-expressing cells, respectively. Scale bars, 10 µm.

Anti-Gαo signal was absent in non-transfected HEK293T cells, consistent with the lack of Gαo expression in the parental HEK293 line^54^. ACh stimulation triggered robust internalization of the M_2_R-YFP-Gγ3 complex in the presence or absence of wild-type Gαo expression, suggesting that endogenous Gα subunits were sufficient to engage the receptor (Fig. 4B). Notably, co-expression of the DEE17/NEDIM variants markedly inhibited both M_2_R phosphorylation and endocytosis, contrasting with the normal response observed in adjacent Gαo-negative cells (Fig. 4C-F).

Finally, we moved beyond the split-YFP assay and analyzed ACh-mediated phosphorylation of GFP-M_2_R upon co-expression of untagged Gαo variants (Fig. 5A). We selected the GFP-M_2_R construct because it lacks intracellular tags, thereby avoiding potential artifacts originating from the receptor. We first examined whether the PM localization of GFP-M_2_R was affected by any Gαo variant. To this end, we immunostained the extracellular M_2_R pool under non-permeabilizing conditions using an anti-GFP antibody, and normalized the signal to the total GFP fluorescence to account for differences in expression (Fig. S8A-H). Quantification showed that GFP-M_2_R surface targeting was unaffected by co-expression of Gαo variants (Fig. S8I).

**Figure 5.**
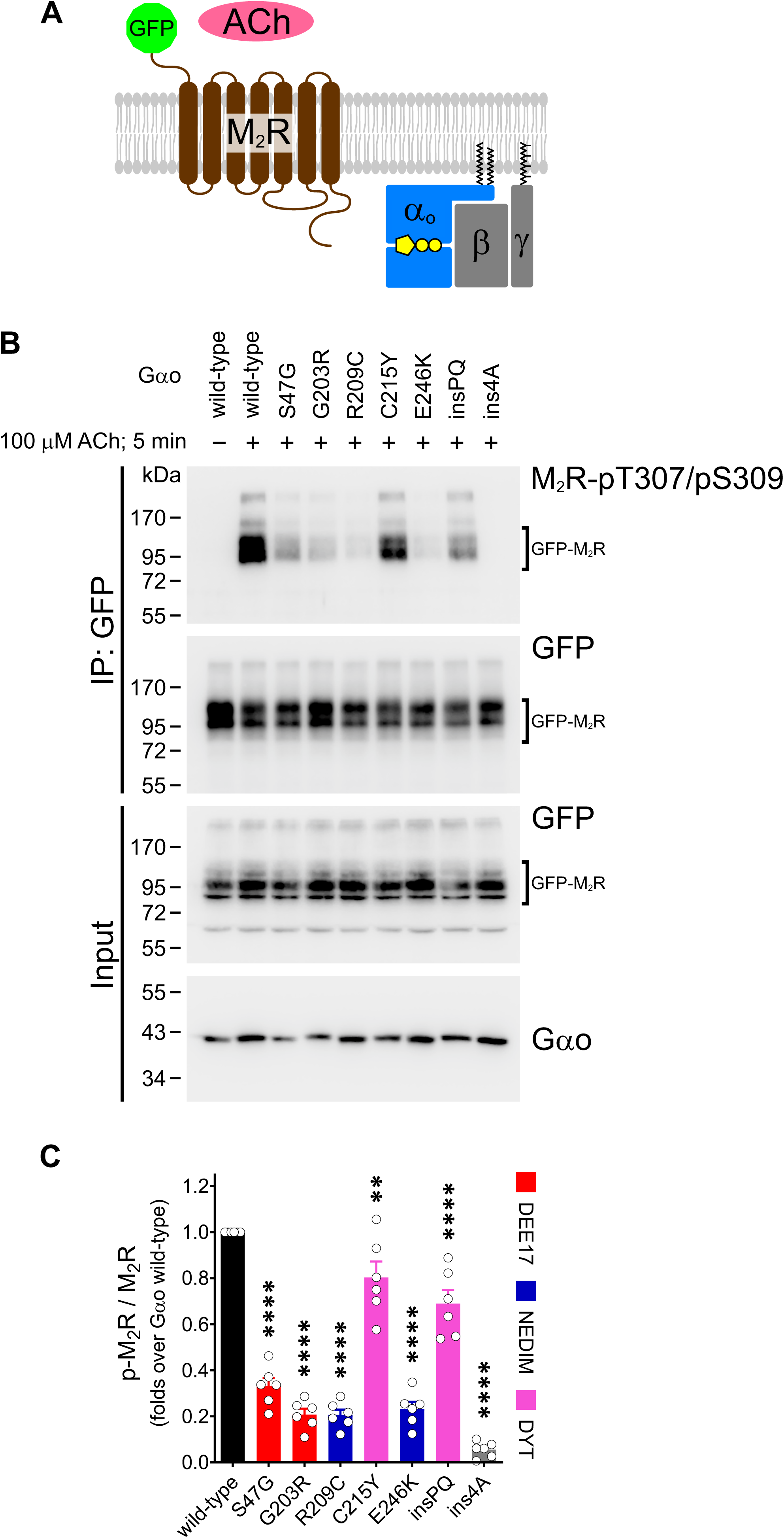
Immunoprecipitation of GFP-M_2_R validates dominant GPCR coupling by pathogenic Gαo variants. (**A**) Diagram of the GFP-M_2_R construct and heterotrimeric Gαoβγ, illustrating the lack of intracellular tagging in the proteins. (**B**) HEK293T cells co-expressing GFP-M_2_R and Gαo wild-type, pathogenic mutants, or the ins4A control were stimulated with 100 µM acetylcholine (ACh) for 5 min and subjected to immunoprecipitation (IP) using an anti-GFP nanobody. Western blotting and immunodetection were performed with antibodies against pT307/pS309 and GFP. (**C**) Quantification of M_2_R phosphorylation levels relative to total receptor (*n* = 6). Gαo mutant associations with DEE17, NEDIM, and DYT phenotypes are color-coded. Data represent mean ± SEM. Statistical analysis was performed using one-way ANOVA followed by Dunnett’s multiple comparisons test; \*\**p* < 0.005 and \*\*\*\**p* < 0.0001.

As expected, IP analysis revealed no detectable phosphorylation of GFP-M_2_R at steady state, whereas a strong anti-pT307/pS309 signal was observed following 5 min of ACh stimulation (Fig. 5B). Remarkably, all pathogenic Gαo mutants reduced GFP-M_2_R phosphorylation, following a pattern closely resembling that observed with the split-YFP assay. Quantification revealed a pronounced 65-80% reduction in M_2_R phosphorylation upon co-expression of the DEE17/NEDIM variants, and ∼95% reduction for the Gαo ins4A mutant (Fig. 5C). In contrast, co-expression of the DYT variants C215Y and insPQ led to more modest reductions of 20% and 30%, respectively.

Altogether, this study indicates that pathogenic Gαo variants remain persistently and dominantly coupled to activated GPCRs. These findings highlight a critical pathogenic mechanism in *GNAO1*-related disorders and establish a robust split-YFP assay for its detection and potential therapeutic targeting.

## Discussion

Pathogenic *GNAO1* mutations disrupt multiple aspects of Gαo function^26–28^. Two distinct mechanisms – nonproductive GPCR coupling and Gβγ sequestration – have been independently reported for several Gαo variants^26–28^. This apparent contradiction, together with the absence of a unified biosensor approach, prompted us to develop a split-YFP assay to directly visualize the dominant GPCR coupling behavior of Gαo mutants. Using this system, we found that persistent, dominant coupling to activated GPCRs is a common feature among pathogenic Gαo variants. The most severe mutants failed to disengage from activated receptors, thereby blocking GRK-mediated phosphorylation and subsequent endocytosis. In contrast, less severe mutants only partially impaired GPCR phosphorylation without preventing internalization.

BiFC assays have been widely used to analyze protein–protein interactions in living cells^33^, including studies on GPCRs, where most applications have focused on receptor homo- and heteromerization^38, 57–59^. Other BiFC-based approaches have visualized Gβγ dimer formation and localization^60^, Src association with GPCRs^38^, and β-arrestin recruitment following receptor activation^61, 62^. To our knowledge, this study represents the first BiFC system specifically designed to monitor GPCR coupling by a Gα subunit in mammalian cells. In this assay, YFP complementation at the PM between GPCRs and the Gα appears to be nonselective, as Gαo was able to efficiently tether to the Gs-coupled β_2_AR. Upon stimulation, however, the assay appears highly specific, since pathogenic Gαo variants disrupted internalization of Gi/o-coupled GPCRs, but had no effect on β_2_AR endocytosis. In addition to its functional specificity, another advantage of this system is that the complemented split-YFP can be readily isolated using an anti-GFP nanobody^55, 63^. This enables not only biochemical analysis of the GPCR-YFP-Gα complex, but also opens possibilities for structural studies of dominant GPCR coupling by pathogenic Gαo variants.

Dominant GPCR coupling by Gα subunits has long been recognized in the GPCR field^51, 52, 64–66^. Studies using engineered, non-pathogenic Gα mutations have proposed two mechanistic modes: sequestration of the activated GPCR by the heterotrimeric G protein, or by nucleotide-free Gα subunits. Although our split-YFP assay cannot discriminate between these mechanisms, insight can be drawn from the distinct behaviors of Gαo variants in previous BRET-based assays. For example, the S47G mutant likely traps the receptor – and Gβγ – in an open, intermediate activation state of the heterotrimeric complex, as it was reported to retain near-normal association with Gβγ but shows markedly impaired dissociation upon GPCR activation^26–28^. In contrast, the recurrent G203R, R209C, and E246K variants exhibit significant Gβγ dissociation upon stimulation^26–28, 67^, consistent with receptor trapping by nucleotide-free Gαo following Gβγ dissociation (Fig. 6). Nevertheless, distinguishing between these two modes will require further mechanistic studies.

**Figure 6.**
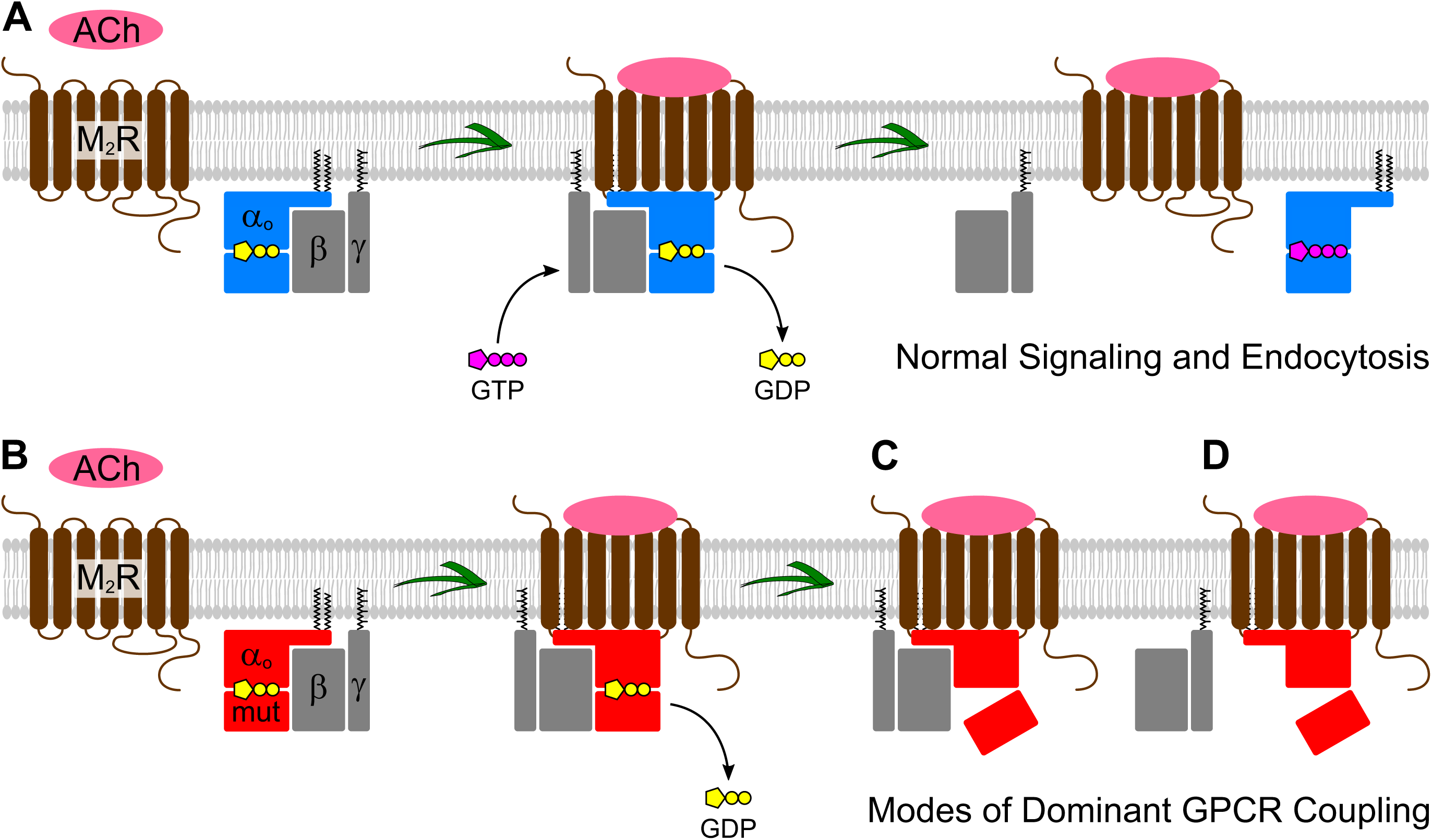
Proposed mechanisms of dominant GPCR coupling by pathogenic Gαo mutants. (**A**) Under physiological conditions, acetylcholine (ACh) activates the M_2_ muscarinic acetylcholine receptor (M₂R), promoting GDP-GTP exchange on wild-type Gαo (blue) within the heterotrimeric G protein. This activation triggers dissociation of Gαo-GTP and Gβγ subunits, initiating downstream signaling and allowing receptor phosphorylation and endocytosis. (**B**-**D**) Pathogenic Gαo variants (red) engage activated GPCRs leading to GDP release (**B**). The failure of Gαo mutants to adopt the active GTP-bound conformation results in dominant GPCR coupling through two distinct mechanisms. In one mode, the mutant Gαo remains in an open, intermediate activation state that retains both the GPCR and Gβγ within a persistent heterotrimeric complex (**C**). In the second mode, mutant Gαo remains tightly bound to the activated receptor in a nucleotide-free state after Gβγ dissociation (**D**). Both mechanisms impair receptor phosphorylation and block endocytosis, potentially leading to signaling defects.

In addition to Gαo, dominant GPCR coupling has thus far been implicated only in pathogenic Gαi3 mutations^68^. However, several residues homologous to the Gαo positions analyzed here – Ser47, Gly203, Arg209, and Glu246 – are also mutated in other Gα subunits linked to rare genetic conditions: Gαi2 G203R and R209W^69^, Gαolf E255A^70^, Gαs R231C/H and E268K/G^71^, and Gαi3 S47R, which was shown to dominantly couple to the endothelin type A receptor^68^. These observations support the possibility that dominant GPCR coupling may extend beyond Gαo and Gαi3, as many disease-associated mutations affect additional highly conserved residues shared across Gα subunits^12–17^. Thus, we envision that this simple split-YFP approach could be adapted for routine testing of Gαo and potentially other pathogenic Gα variants implicated in a growing number of rare genetic disorders.

Beyond mechanistic insights, the robustness and scalability of the split-YFP system lay a foundation for drug discovery or repositioning efforts aimed at restoring normal GPCR coupling dynamics. Such applications could accelerate the identification of therapeutic compounds that reverse receptor trapping or promote dissociation of Gαo mutants, offering new avenues for targeted therapies in *GNAO1*-related disorders.

## Methods

### Antibodies and reagents

Primary antibodies (Abs) for immunofluorescence (IF) and Western blots (WBs): monoclonal Abs (mAbs) anti-Gαo (clone A2, sc-13532; IF: 1:50) and anti-Gαo (clone E1, sc-393874; WB: 1:250) were from Santa Cruz Biotechnology; mAb anti-HA-tag (11867423001; IF: 1/500) was from Roche. Polyclonal antibodies (pAbs) anti-M_2_R (7TM0014N; WB: 1/1000) and anti- pT307/pS309 (7TM0014D; IF: 1:500, WB: 1:1000) were from 7TM Antibodies; pAbs anti-GFP (PABG1; IF: 1/1000) and anti-GFP (50430-2-AP; WB: 1:5000) were from Proteintech.

All secondary Abs for immunofluorescence (IF) and Western blots (WBs) were from Jackson ImmunoResearch: anti-Mouse Cy5-conjugated (715-175-151; IF: 1/500), anti-Rabbit Cy3- conjugated (711-165-152; IF: 1/500), anti-Rat Cy3-conjugated (112-165-143; IF: 1/500), anti-mouse HRP-conjugated (115-035-146; WB: 1:5000), and anti-rabbit HRP-conjugated (111-035-144; WB: 1:5000).

Acetylcholine (A2661), isoproterenol (I6504), and DAPI (32670) were from Sigma-Aldrich, and EGF-Rhodamine (E3481) was from Thermo Fisher Scientific. Fentanyl (CAS no. 1443- 54-5) were obtained from the University Hospital of Geneva with Département de la sécurité, de la population et de la santé (DSPS) authorization from the Canton of Geneva.

### Plasmids and molecular cloning

Plasmids encoding for the untagged Gαo wild-type and pathogenic mutants were published earlier^28, 31, 41^. Untagged Gαo ins4A was kindly provided by Nevin A. Lambert (Augusta University, GA, USA). To generate the mRFP internal tagging of Gαo (at Gly92; Gαo-mRFP), the mRFP sequence was amplified by PCR from pmRFP-C1^41^ using the oligonucleotide primers For: 5’–GTCGCCGGGCCCGCCTCCTCCGAGGAC–3’ and Rev: 5’–TTTAAAGCAAGTAAAACCTC–3’. The PCR-product was cloned in-frame into the PspOMI/BsrGI sites of Gαo-GFP^63^. The sequences for the split-YFP fragments YN (aa 1- 158) and YC (aa 159-239) containing a flexible 10-amino acid (GGGGSGGGGS) linker were kindly provided by Daniel F. Legler (Institute of Cell Biology and Immunology Thurgau, Switzerland). To produce the Gαo-YC constructs (internal YC insertion at Gly92), the YC sequence was amplified by PCR using the primers For: 5’– GGATCCACCGGTCCTCACCGGGCCCGGTGGCGGTGGCTCTGG–3’ and Rev: 5’–AGATCTGAGTCCGGACTTGTACACGGACCC–3’. The resulting product was cloned in-frame into the PspOMI/BsrGI sites of Gαo-GFP wild-type and mutants^28, 31^. The Gαo-YC for ins4A was obtained by replacing the AgeI/PspOMI fragment in Gαo-YC wild-type with the AgeI/NotI fragment from untagged Gαo ins4A. The GFP-M_2_R construct was generated by cloning in-frame the fragment blunted-EcoRI/XhoI from the untagged M_2_R plasmid (MAR0200000; cDNA Resource Center) into the Eco53KI/SalI sites of the PrP-leader-GFP plasmid^72^. To produce the M_2_R-YN construct, we first PCR-amplified the YN sequence with the primers For: 5’–AGCCCGGACCGGTCCTCACCATGGATGGTGGCGGTGGCTC–3’ and Rev: 5’–TTTAGGTGGCGGCCGCGAATAGGACCCTCTAGATTAC–3’. The PCR-product was digested with AgeI/NotI and ligated in-frame into the AgeI/NotI sites of M_2_R-NLuc^28^. The sequence for a signal peptide-HA-tag was generated by the primers For: 5’– TCGACCACCATGAAGACGATCATCGCCCTGAGCTACATCTTCTGCCTGGTATTCGCCTACCCATACGATGTTCCTGACTATGCGG–3’ and Rev: 5’– AATTCCGCATAGTCAGGAACATCGTATGGGTAGGCGAATACCAGGCAGAAGATGTAGCTCAGGGCGATGATCGTCTTCATGGTGG–3’ and cloned in-frame into the XhoI/EcoRI sites upstream of the M_2_R. For the M_2_R-YC plasmid, the YC fragment was amplified by PCR using the primers For: 5’–ACTATGAACCGGTACTCACCATGGATGGTGGCGGTGGCTC–3’ and Rev: 5’–GCAACTAGAAGGCACAGTCGAGG–3’, and cloned in-frame into the AgeI/NotI sites of the final M_2_R-YN. The β_2_AR-YN plasmid was obtained by cutting the NheI/AgeI coding fragment from β_2_AR-mCFP^73^ (Addgene plasmid 38260) and insertion into the same sites of M_2_R-YN. The MOR-YN construct was produced by cutting the XhoI/XmaI coding sequence from the MOR-GFP plasmid^74^ (provided by Miriam Stoeber; University of Geneva, Switzerland), and ligation into the XhoI/AgeI sites of M_2_R-YN. The T2A-based bicistronic Gβ3-YN-Gγ3 construct – referred to as YN-Gγ3 for simplicity – was generated through a two-step cloning. First, the single YN-Gγ3 plasmid was cloned by PCR amplification of the YN sequence using the primers For: 5’– CTATAGGACCGGTCAAGACCATGGTGAGCAAGGGCGAGGA–3’ and Rev: 5’– TGCTCGAGTGTACACGGACCCACCACCTCCAGAGC–3’, and the YN fragment was cloned in-frame into the AgeI/BsrGI sites of GFP-Gγ3^41^. Then, a T2A-based bicistronic Gβ3- cpVenus-Gγ9 plasmid was created by cutting the NheI/SalI insert from Go1-CASE^75^ (provided by Gunnar Schulte; Karolinska Institutet, Sweden) and ligation of the fragment into the same sites of pEGFP-C1 (Clontech). The final YN-Gγ3 construct was generated by cutting the YN-Gγ3 coding sequence with AfeI/PspOMI, and ligation of this fragment into the blunted-AgeI/PspOMI of Gβ3-cpVenus-Gγ9. A T2A-based bicistronic untagged Gβ3-Gγ9 plasmid (referred to as Gβ3γ9) was generated by cutting out the cpVenus sequence from Gβ3-cpVenus-Gγ9 with AgeI/BspEI, and religation to produce an in-frame Gβ3-T2A-Gγ9 construct.

### Cell lines and culture conditions

Human HEK293T (CRL-3216, ATCC) cells were grown in DMEM, supplemented with 10% FCS, 2 mM L-glutamine, and 1% penicillin-streptomycin (all from Thermo Fisher Scientific) at 37°C and 5% CO_2_. All vector transfections were carried out with X-tremeGENE HP (XTGHP-RO, Roche) according to the manufacturer’s instructions.

### Split-YFP-based BiFC assay and microscopy

For the split-YFP assay between GPCRs and Gαo, HEK293T cells were co-transfected with GPCR-YN constructs (M_2_R, MOR, or β_2_AR), Gαo-YC (wild-type, pathogenic mutants, or the control ins4A), and Gβ3γ9 at a plasmid ratio of 2.5:2.5:1. For the split-YFP assay between M_2_R and Gγ3, cells were co-transfected with M_2_R-YC, YN-Gγ3, and untagged Gαo variants or empty plasmid at the same ratio. After for 6 h, cells were trypsinized, seeded onto poly-L-lysine-coated coverslips, and cultured – or co-cultured, in the case of the M_2_R-Gγ3 split-YFP assay – for an additional 15 h in complete DMEM.

For endocytosis analysis of GPCR-YFP-Gαo complexes, cells were stimulated for 10 min with 100 µM acetylcholine (ACh) for M_2_R, 1 µM fentanyl for MOR, or 10 µM isoproterenol for β_2_AR. To determine M_2_R phosphorylation, cells were stimulated with 100 µM ACh for 7 min. To evaluate EGF-uptake, cells were treated with 10 ng/ml EGF-Rhodamine for 10 min as previously described^46^.

After treatments, cells were fixed with 4% PFA in PBS for 20 min and permeabilized with ice-cold 0.1% Triton X-100 in PBS for 1 min – this step was omitted under non-permeabilizing conditions. Cells were then blocked for 1 h in PBS supplemented with 1% BSA (810533; Millipore), incubated with primary antibodies in blocking buffer for 2 h at room temperature (RT), washed, and subsequently incubated with fluorescently labelled secondary antibodies and DAPI in blocking buffer for 2 h at RT. Coverslips were mounted with VECTASHIELD (H- 1700; Vector Laboratories) on microscope slides. For immunostaining against phosphorylated M_2_R, all steps were performed in TBS instead of PBS.

Samples were recorded using a Plan-Apochromat 63x/1.4 oil objective on an LSM800 confocal microscope with ZEN 3.7 software (all from Zeiss). When quantification was required, all images were acquired using identical confocal settings across all channels. Mean fluorescence intensities, cell area, and vesicle-like structure counts were determined from confocal images either manually or semi-automatically using in-house scripts, all with ImageJ v1.54p (National Institutes of Health).

### PM targeting of GFP-M_2_R

HEK293T cells were co-transfected with GFP-M_2_R, untagged Gαo variants (wild-type, pathogenic mutants, or the control ins4A), and Gβ3γ9 at a plasmid ratio of 2.5:2.5:1. Samples were immunostained under non-permeabilizing conditions, and image acquisition and quantification were performed as described above.

### Immunoprecipitation

The recombinant GST-tagged nanobody against GFP^55^ was expressed in *Escherichia coli* Rosettagami (71351; Novagen) and purified using Glutathione Sepharose 4B beads according to the manufacturer’s instructions. Protein purity was confirmed by SDS-PAGE and Coomassie blue staining.

HEK293T cells were co-transfected with the split-YFP constructs for M_2_R, Gαo, and Gβ3γ9, or with GFP-M_2_R, untagged Gαo variants, and Gβ3γ9, using the same plasmid ratio described above. After 24 h of transfection, cells were stimulated with 100 µM ACh for 5 min, then immediately resuspended in ice-cold TBS-lysis buffer (20 mM Tris-HCl, pH 7.5, 150 mM NaCl, 1% Triton X-100, 10% glycerol) supplemented with protease (04693132001) and phosphatase (04906837001) inhibitor cocktails from Roche. Lysates were passed 15 times through a 25-G needle and cleared by centrifugation at 15,000xg for 15 min at 4°C. Supernatants were incubated with 2 μg of purified anti-GFP nanobody for 30 min on ice, followed by addition of 20 μL of Glutathione Sepharose 4B beads (17075601; Cytiva) and rotation for 3 h at 4°C.

Beads were washed repeatedly with TBS-lysis buffer and resuspended in Laemmli sample buffer. Inputs and beads were heated at 50°C for 30 min and analyzed by SDS-PAGE Western blotting using Abs against phospho-M_2_R (pT307/pS309), total M_2_R, Gαo, and GFP. HRP-conjugated secondary antibodies were used for detection, with all antibody incubations performed in TBS containing 1% BSA. Signal was visualized using enhanced chemiluminescence (ECL) on a Fusion FX6 Edge system (Vilber). Quantification of all WBs was performed using ImageJ v1.54p. Final image preparation was done in EvolutionCapt v18.11 (Vilber) and CorelDRAW 2020.

## Data availability

All data supporting the findings of the study are included within the article.

## Acknowledgements

This study was supported by Famiglie GNAO1 Association grant to GPS. We are very thankful to Arthur Radoux-Mergault for the valuable discussions and insightful input throughout the development of this project. We thank Sabina Troccaz for her excellent technical assistance. We are grateful to all members of the Bioimaging core facility at the CMU for their expert assistance with microscopy.

---

Figure S1. **Subcellular localization of M_2_R and Gαo in HEK293T cells**

(**A**,**B**) Confocal images of HEK293T cells expressing N-terminally GFP-tagged M_2_R (GFP-M_2_R; **A**) and Gαo internally tagged with mRFP (Gαo-mRFP; **B**). GFP-M_2_R-expressing cells were immunostained with anti-GFP under non-permeabilizing conditions to label surface GFP. Boxed regions are shown at higher magnification in the lower panels. Scale bars, 10 µm.

Figure S2. **ACh-mediated Stimulation HEK293T cells expressing the M_2_R-YFP-Gαo complex**

(**A**-**G**) Representative confocal images of HEK293T cells expressing the M_2_R-YFP-Gαo complex containing the indicated Gαo variants. Cells were immunostained under non-permeabilizing conditions to label surface M_2_R at steady state (left panels) and after 10 min of acetylcholine (ACh) stimulation (right panels). Scale bars, 10 µm. (**H**) Quantification of M_2_R-YFP-Gαo complex formation across different Gαo variants. Gαo mutant associations with DEE17, NEDIM and DYT phenotypes are color-coded. Box plot indicate median (middle line), 25th, 75th percentile (box), and lowest, highest value (whiskers); two-three independent experiments (wild-type, *n* = 78; S47G, *n* = 55; G203R, *n* = 59; R209C, *n* = 78; C215Y, *n* = 59; E246K, *n* = 63; insPQ, *n* = 54). (**I**) HEK293T cells expressing the split-YFP Gαo-YC construct (wild-type and pathogenic mutants) were analyzed by Western blot using an anti-Gαo antibody. (**J**) Quantification of Gαo-YC mutant expression levels relative to wild-type (*n* = 3). Data represent mean ± SEM. Statistical analyses were performed using one-way ANOVA followed by Dunnett’s multiple comparisons test; \**p* < 0.05, \*\*\**p* < 0.001, \*\*\*\**p* < 0.0001, and ns: not significant.

Figure S3. **EGF-Rhodamine uptake by HEK293T cells expressing the M_2_R-YFP-Gαo complex**

(**A**-**G**) Representative confocal images of HEK293T cells expressing the M_2_R-YFP-Gαo complex with the indicated Gαo variants. Cells were incubated with 10 ng/ml of EGF-Rhodamine for 10 min. Nuclei were visualized by DAPI staining. Cyan lines demarcate cells expressing M_2_R-YFP-Gαo. Scale bars, 10 µm. (**H**) Quantification of EGF-Rhodamine uptake. Gαo mutant associations with DEE17, NEDIM and DYT phenotypes are color-coded. Box plot indicate median (middle line), 25th, 75th percentile (box), and lowest, highest value (whiskers); two independent experiments (wild-type, *n* = 51; S47G, *n* = 56; G203R, *n* = 51; R209C, *n* = 52; C215Y, *n* = 57; E246K, *n* = 52; insPQ, *n* = 55). Statistical analysis was done using one-way ANOVA followed by Dunnett’s multiple comparisons test; ns: not significant.

Figure S4. **Clinically severe Gαo mutants disrupt µ-opioid receptor (MOR) endocytosis**

(**A**) Illustration of the split-YFP assay applied to MOR and heterotrimeric Gαoβγ. (**B**,**C**) Representative confocal images of HEK293T cells expressing the MOR-YFP-Gαo complex at steady state (**B**) and after 10 min of 1 µM fentanyl stimulation (**C**). (**D**-**I**) Confocal images of fentanyl-stimulated HEK293T cells expressing the MOR-YFP-Gαo complex with the indicated pathogenic Gαo variants. Scale bars, 10 µm.

Figure S5. **Pathogenic Gαo variants do not affect β_2_-adrenoceptor (β_2_AR) endocytosis**

(**A**) Depiction of the split-YFP assay applied to β_2_AR and heterotrimeric Gαoβγ. (**B**,**C**) Representative confocal images of HEK293T cells expressing the β_2_AR-YFP-Gαo complex at steady state (**B**) and after 10 min of 10 µM isoproterenol stimulation (**C**). (**D**-**I**) Confocal images of isoproterenol-stimulated HEK293T cells expressing the β_2_AR-YFP-Gαo complex assembled with the indicated Gαo mutants. Scale bars, 10 µm.

Figure S6. **The non-pathogenic Gαo ins4A construct blocks M_2_R and MOR endocytosis**

(**A**-**F**) Representative confocal images of HEK293T cells expressing the M_2_R-YFP-Gαo (**A**- **C**) and MOR-YFP-Gαo (**D**-**F**) complexes assembled with either Gαo wild-type (**A**,**B**,**D**,**E**) and the non-pathogenic ins4A variant (**C**,**F**). Cells were analyzed at steady state (**A**,**D**) and after 10 min stimulation with 100 µM acetylcholine (ACh; **B**,**C**) or 1 µM fentanyl (**E**,**F**). Scale bars, 10 µm.

Figure S7. **Immunoprecipitation of the M_2_R-YFP-Gαo complex**

(**A**) HEK293T cells expressing the split-YFP constructs M_2_R-YN and Gαo-YC, either individually or in combination as indicated, were subjected to immunoprecipitation (IP) using an anti-GFP nanobody. Western blotting and immunodetection were performed with antibodies against M_2_R and Gαo. Arrowheads indicate M_2_R-YN, Gαo-YC, and the monomeric and dimeric forms of the M_2_R-YFP-Gαo complex.

Figure S8. **Plasma membrane localization of GFP-M_2_R in HEK293T cells**

(**A**-**H**) Confocal images of HEK293T cells expressing N-terminally GFP-tagged M_2_R (GFP-M_2_R) along with untagged Gαo variants, as indicated. Cells were immunostained with anti-GFP under non-permeabilizing conditions to label extracellular GFP, and counterstained with DAPI to visualize nuclei. Scale bars, 10 µm. (**I**) Quantification of surface M_2_R levels, measured as the ratio of surface to total GFP-M_2_R signal. Gαo mutant associations with DEE17, NEDIM and DYT phenotypes are color-coded. Box plot indicate median (middle line), 25th, 75th percentile (box), and lowest, highest value (whiskers); two independent experiments (wild-type, *n* = 66; S47G, *n* = 67; G203R, *n* = 60; R209C, *n* = 63; C215Y, *n* = 69; E246K, *n* = 64; insPQ, *n* = 65; ins4A, *n* = 64). Statistical analysis was done using one-way ANOVA followed by Dunnett’s multiple comparisons test; ns: not significant.

