## Supplemental Figures 1 to 8 for "Visualizing the Dominant GPCR Coupling of Pathogenic Gαo Mutants in *GNAO1*-Related Disorders"

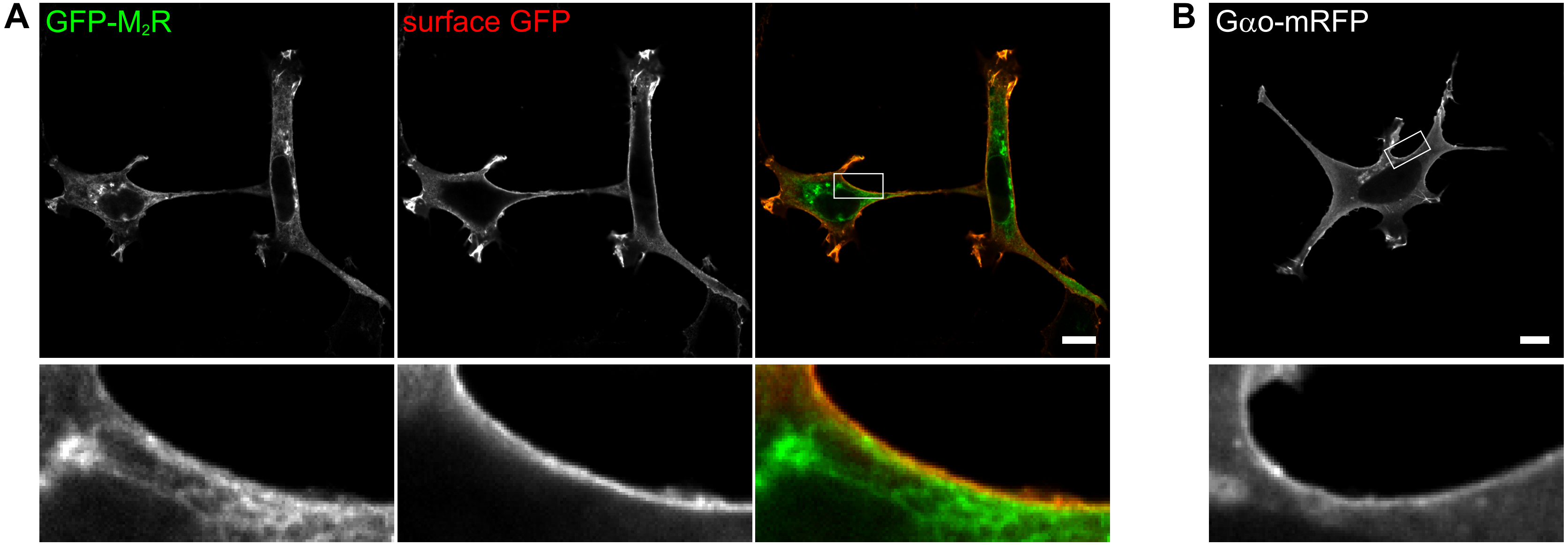


Larasati et al. Supp Fig. S1

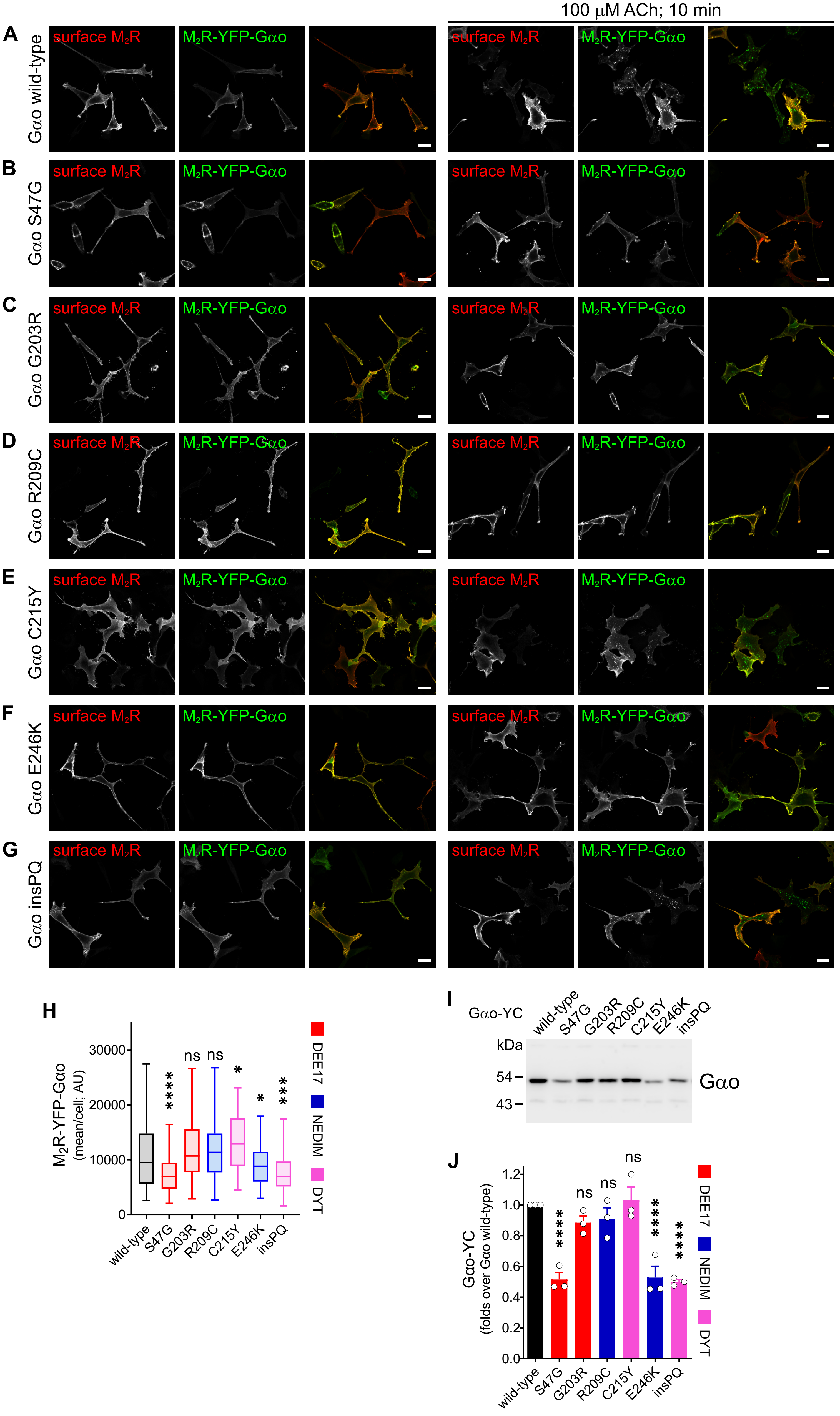


Larasati et al. Supp Fig. S2

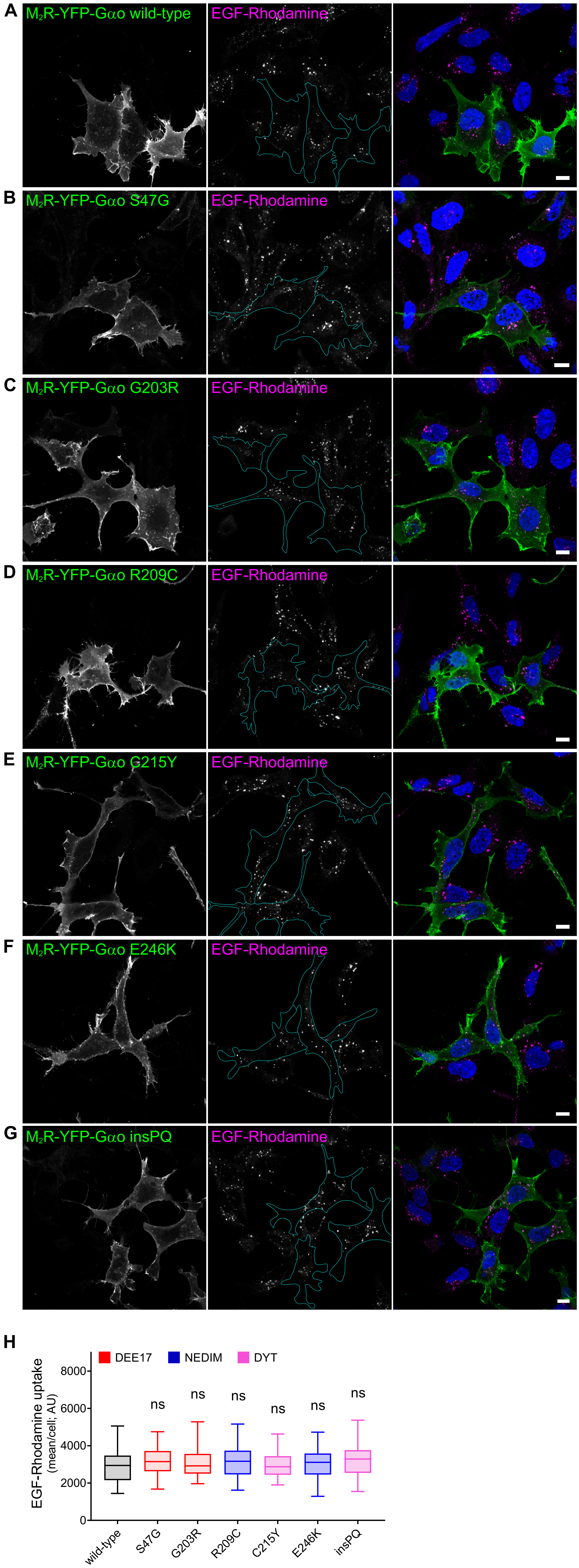


Larasati et al. Supp Fig. S3

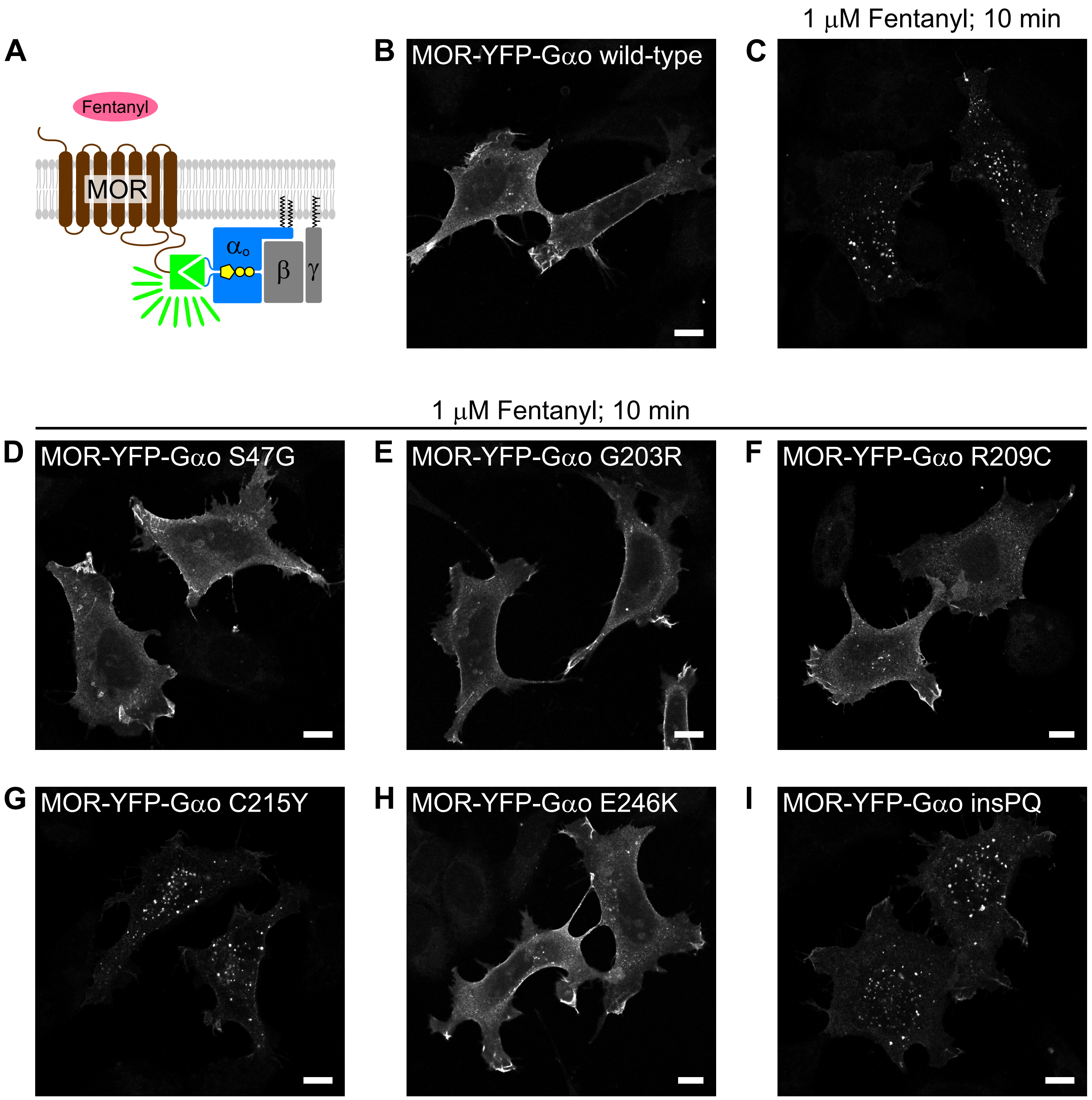


Larasati et al. Supp Fig. S4

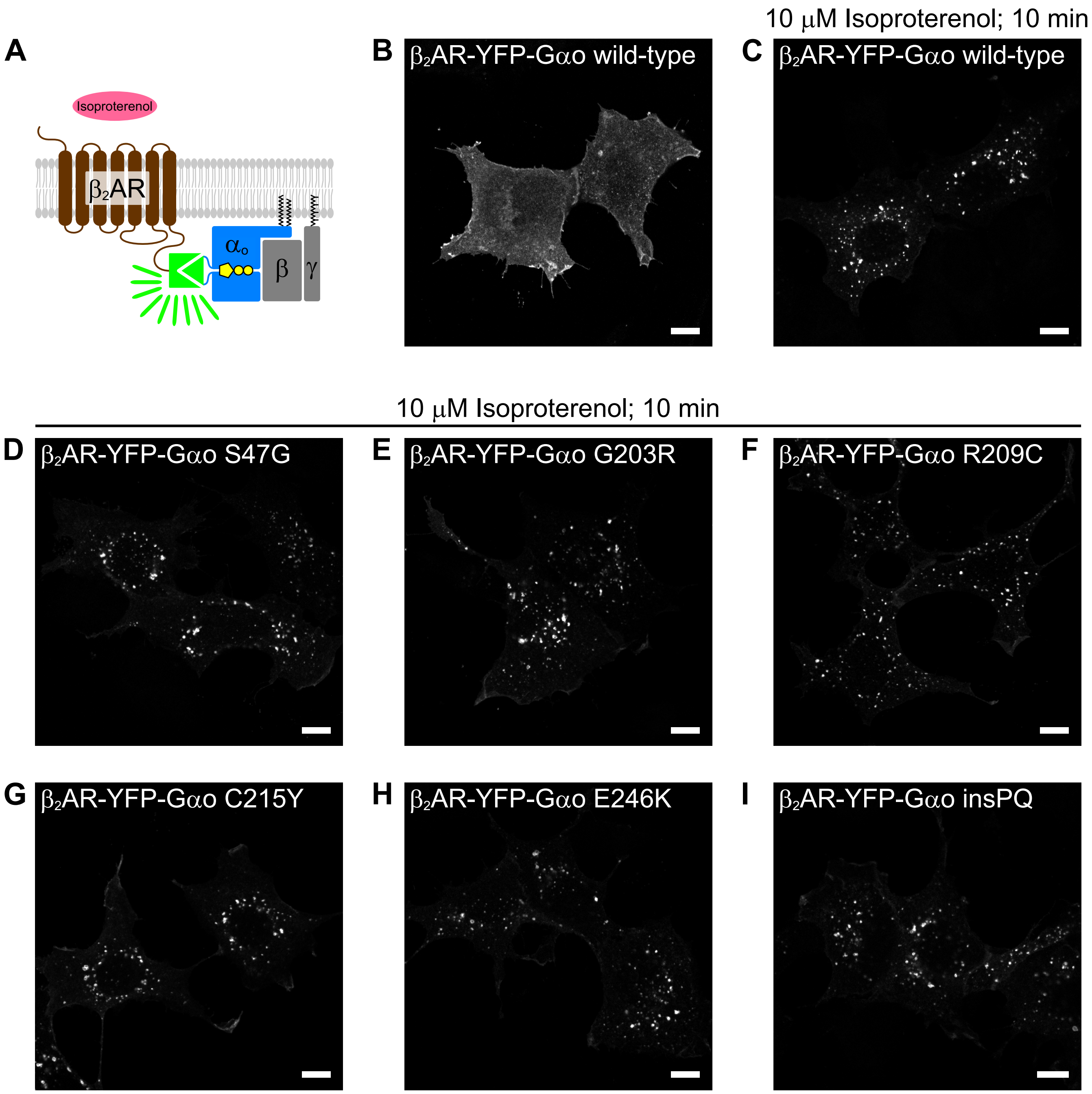


Larasati et al. Supp Fig. S5

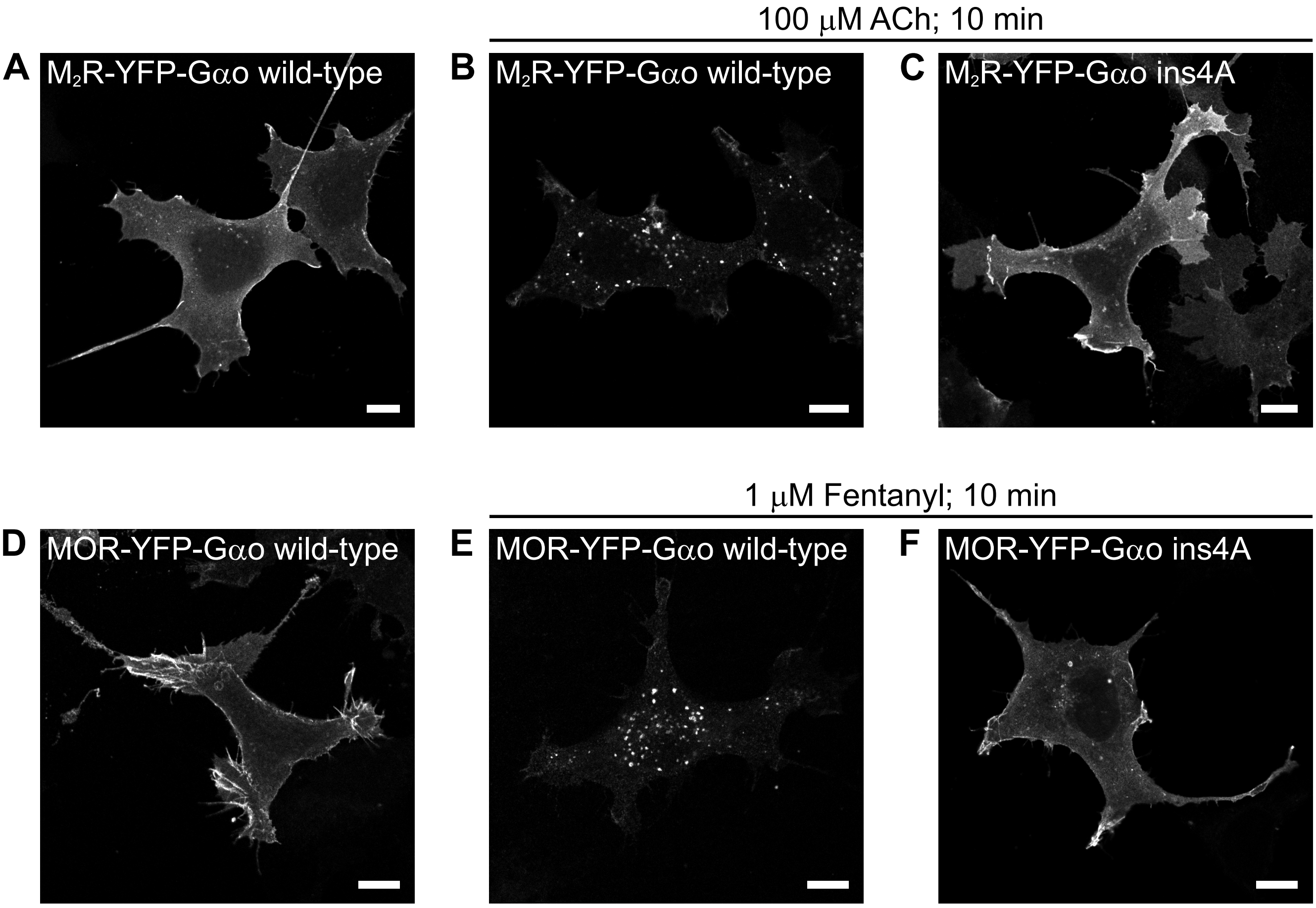


Larasati et al. Supp Fig. S6

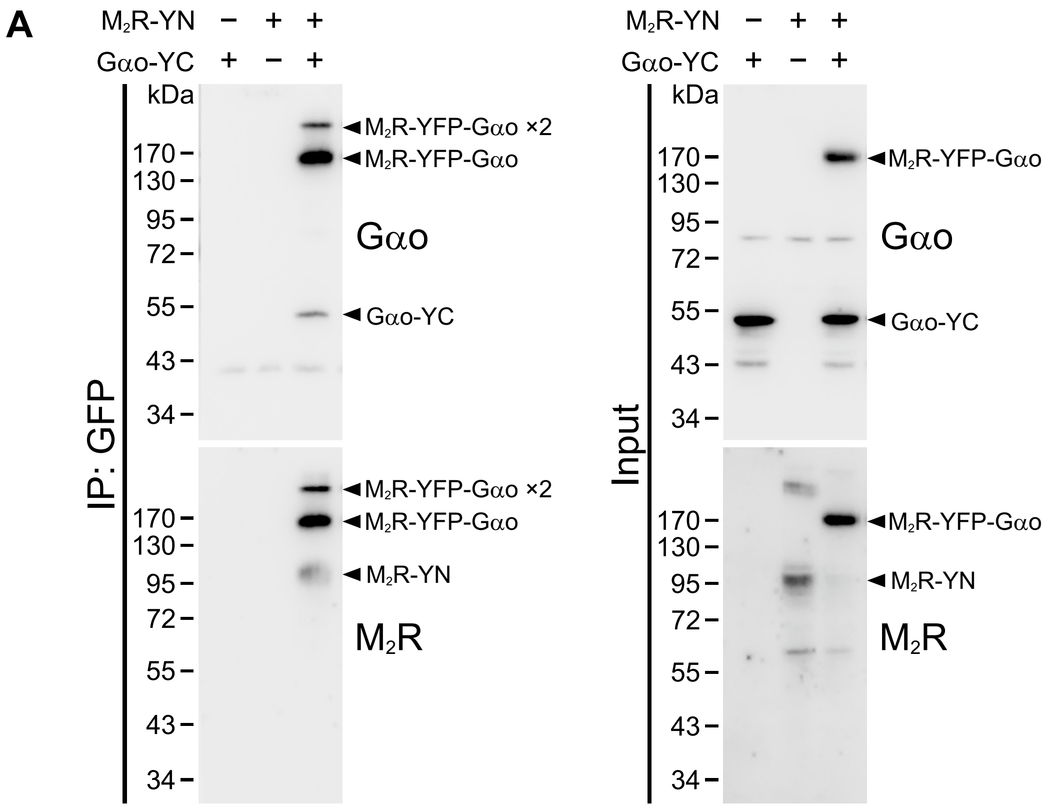


Larasati et al. Supp Fig. S7

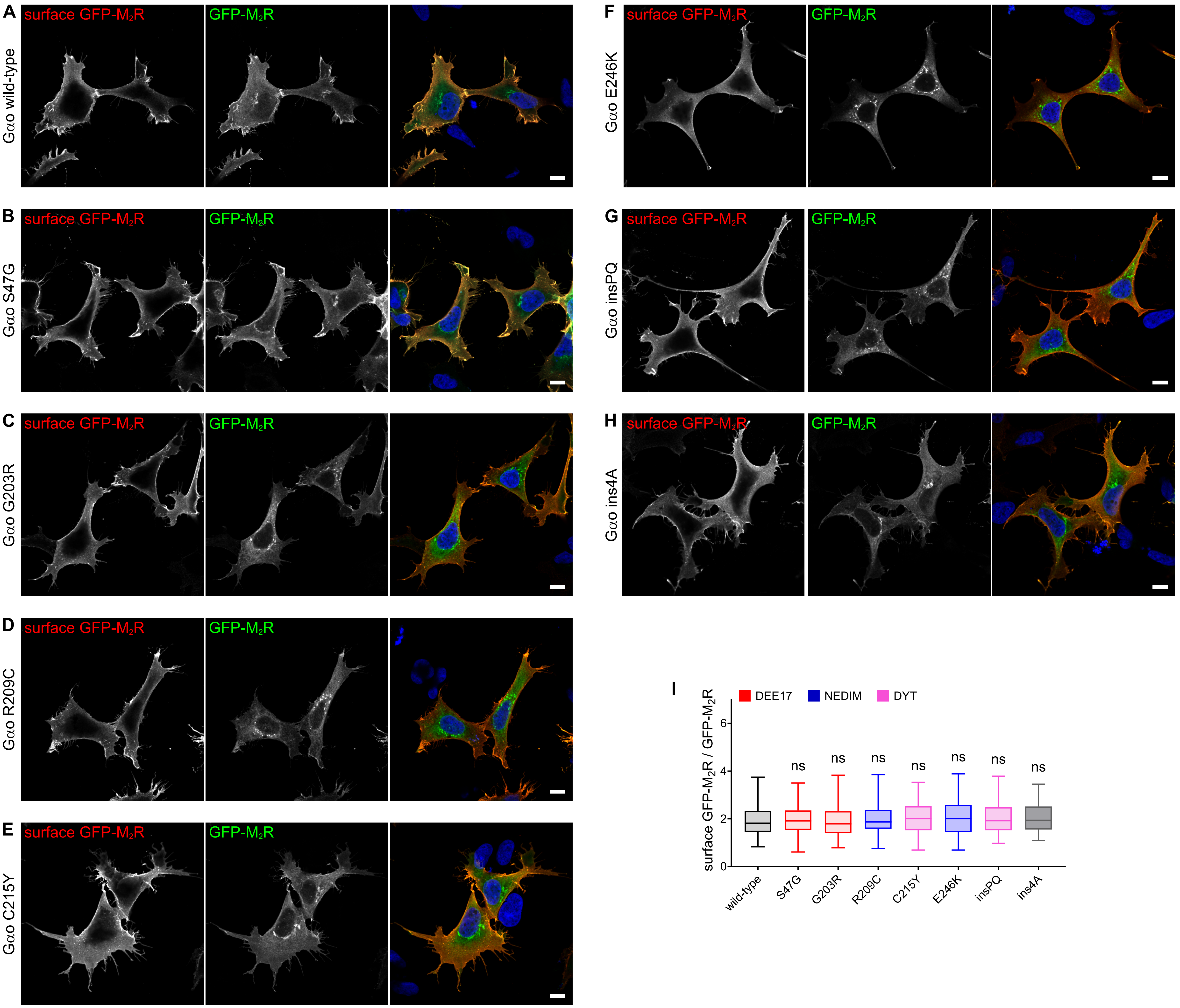


Larasati et al. Supp Fig. S8
